# Transcription Factor HNF4A is Involved in Breast Cancer Recurrence

**DOI:** 10.1101/2024.03.23.586428

**Authors:** Rahul Valiya Veettil, Noah Eyal Altman, Ruth Hashkes, Eitan Rubin

**Author notes:** **Correspondence:** Eitan Rubin.

## Abstract

Recurrence is a major challenge for breast cancer (BC) management, yet its mechanism is poorly understood. Although computational analysis of expression compendia has proved useful for studying BC recurrence, some recently developed methods have not been used for this purpose. Here we present a bioinformatics reanalysis of a compendium containing 1519 cases with documentation of expression and information about recurrence with >5y follow-up. A compendium of expression profiles were divided into sub-cohorts by time to recurrence, and genes were ranked by the relative significance of expression differences. The top 1% of the genes (131 genes) were used in enrichment and induced network module analyses. The findings were validated in two independent cohorts, E-MTAB-365 and GSE20685. HNF4A was found to be the major hub in the genes whose expression differs most strikingly between short and long/no recurrence patients: it is highly connected to differentially expressed genes, and high expression of HNF4A is associated with longer recurrence-free survival in the compendium, even when it was split by ER status. These results were further validated in two independent cohorts of ER+ + endocrine + chemotherapy-treated patients and found to be concordant. Here we re-analyze a previously published compendium and find that HNF4A is pivotal to expression differences between early and late/non-recurring BC patients. HNF4A has been reported to be associated with cancer risk in several cancers and is speculated to be recurrence-associated in BC. However, we establish for the first time an association between BC time-to-recurrence and HNF4A expression. Further research is required to check if HNF4A expression is causative or just correlative to recurrence. Yet since linoleic acid is a ligand of HNF4A, our finding may explain why consumption of linoleic acid was proposed to affect recurrence risk in BC.

## 1 Introduction

Breast cancer (BC) is the leading cause of cancer-related mortality in women (MP *et al*., 2007). Primary BC is manageable, as it affects non-vital tissue, and it can be treated with surgery and irradiation with relative ease and minimal damage. However, tumor recurrence, especially distant recurrence is too often not manageable: the new tumors may occur as metastases in vital tissues and are typically more resistant to treatment (Coley, 2008). As a result, recurrence is the leading cause of BC mortality.

BC recurs relatively early (1-2 years after surgery) in most cases (Saphner, Tormey and Gray, 1996; Demicheli *et al*., 2004). However, apparently cured BC tumors may recur as late as 10 or even 20 years after surgery (Fisher *et al*., 2002). Tumor cells have been proposed to have already disseminated to distant loci even before the initial diagnosis. In accordance with this hypothesis, bone marrow specimens were found to contain detectable residual cancer cells in up to 40% of non-symptomatic post-surgery BC patients (Braun *et al*., 2003). Recurrence thus represents a major challenge for BC management.

Considering the clinical importance of recurrence, surprisingly little is known about the molecular mechanisms behind it. In BC patients, recurrence-free survival (RFS) was long predicted solely based on tumor size and the extent of lymph node involvement (Valagussa, Tancini and Bonadonna, 1987; Carter, Allen and Henson, 1989). Some prognostic markers have been reported since (van’t Veer and Bernards, 2008), but none play a demonstrated causal role in breast cancer recurrence: the mechanistic basis of recurrence remains speculative. For a long time, recurrence was thought to involve a hypothetical subpopulation of cells that had acquired additional traits, above and beyond malignancy, thus making them suitable for the recurrence process. However, evidence has emerged that challenges this model. Most importantly, expression profiles based on RNA extracted from the primary tumor have been shown to accurately predict recurrence risk (van’t Veer and Bernards, 2008). Since these assays consider expression in bulk, recurrence risk is predicted from gene expression in the majority of the tumor cells and not in a rare subpopulation. As a result, a better understanding of pathway status in tumors that are more likely to recur may shed light on the recurrence mechanism.

One common approach to understanding the pathways and components behind clinical outcomes has been to investigate gene expression. In the context of BC recurrence, this means comparing the expression of genes in primary tumors of patients with early recurrence with gene expression in patients who have shown no recurrence or late recurrence. Subsequently, genes that differ in expression between the groups are identified using statistical methods such as Student’s t-test (Jafari and Azuaje, 2006). Finally, pathways or biological functions enriched in differentially expressed genes are identified (Chen *et al*., 2013; Kuleshov *et al*., 2016). This kind of analysis points to specific pathways or processes that tend to differ between recurring and non-recurrence patients.

Our study applies a modified t-test based approach to rank and choose genes that greatly differ in expression between fast-recurring and late/no recurrence cases. Coupling this approach with the Induced Network Analysis approach, genes that are pivotal to the differences in expression associated with the fast recurrence of BC are identified. We identify HNF4A as pivotal to the expression differences associated with RFS length and show that its expression, as well as the expression of its targets, is associated with time to recurrence. Finally, we propose that since HNF4A is a nuclear receptor of linoleic acid (Yuan *et al*., 2009), it can mark a novel therapeutic approach.

## 2 Materials and Methods

### 2.1 Microarray data: source and preprocessing

The normalized gene expression data and RFS information of 1,809 breast cancer patients were downloaded from the KMplotter website, under supplementary material. For a complete description of the normalization procedures originally used to create these data, see Györffy et. al (Györffy *et al*., 2010; Lánczky *et al*., 2016). Briefly, the authors used raw CEL files, and MAS5 normalized using affy (RRID:SCR_012835) (Irizarry, Gautier and Cope, 2003), Then, probes were chosen only if measured on both GPL96 and GPL570 (n = 22,277), and a second scaling normalization followed setting the average expression on each chip to 1,000 to avoid batch effects. From the original dataset, patients were included only if 5 years or more of follow-up data was available, and no clinical variables missing, leaving 1519 patients for whom 22,215 probes were measured (after removing baseline genes). For the purpose of this work, the resulting matrix is called the “Györffy” dataset. For information about probe selection, see the statistical analysis section below.

### 2.2 Statistical analysis

All statistical analysis was performed in R Project for Statistical Computing (RRID:SCR_001905) (version 3.4.4) (Dessau and Pipper, 2008). For t-statistic and student’s t test, we used 5-year recurrence as the patients’ class and RFS as the independent variable. The t-statistic for every probe was calculated from the difference in mean expression between patients with quick (<5y) or slow/no recurrence (recurrent free for 5y or more). As the degrees of freedom for each of the probes are the same, we consider the absolute t-statistic rather than the p-value for some analyses. The test was run on each probe separately, and for genes with multiple probes, the probe with the most significant p-value was chosen. The 22,215 probes were mapped to 13126 genes using g:Profiler (RRID:SCR_006809) (Reimand *et al*., 2016). The LIMMA (RRID:SCR_010943) (Ritchie *et al*., 2015) was further used to validate the statistical significance of the genes under investigation compared to t-test.

The distribution of t-test statistics was analyzed using mixtool R package (Benaglia *et al*., 2010). Survival analysis and generating Kaplan-Meier plots were performed using survival (RRID:SCR_021137) (Therneau, 2020), and survplot (version 0.0.7; http://www.cbs.dtu.dk/~eklund/survplot/) R packages. ggplot2 (RRID:SCR_014601) (version 0.9.3) (H, 2016), cowplot (RRID:SCR_018081) (Wilke, 2020), and pheatmap (RRID:SCR_016418) (Kolde, 2019) R packages were used for generating density plots and the heatmap. Multiple correction was performed by calculating a t-test p-value cut-off using the compute.FDR function in brainwaver (RRID:SCR_009540) (FDR < =0.01) (version: 1.6, https://cran.r-project.org/src/contrib/Archive/brainwaver/).

### 2.3 Gene signature enrichment analysis

A ranked gene list associated with RFS was created by choosing an arbitrary cutoff of the top 1% of all genes when ranked by t-test p-value (i.e, resulting in 131 significant genes). For overrepresentation analysis, EnrichR (Chen *et al*., 2013; Kuleshov *et al*., 2016) was used with the top 131 genes list as input.

### 2.4 Induced network module analysis

The top 131 ranked genes list was further used to look for induced network modules using CPDB’s induced network module. This tool is described in (Kamburov *et al*., 2009). Briefly, it begins with a seed group of user-provided genes. The tool searches for all the interactions among the seed genes to define a connected network. It also extends the group to infer regulators or interactors which are not present in the seed gene list. To infer intermediate genes, the tool tests the significance of their connectivity to the seed gens group, using a binomial proportion test for each network and deriving a z-score for each inferred gene. An intermediate is added to the network only if the z-score of its connectivity is larger than a user-defined cutoff (we use the default value of 20). This approach uncovers intermediate nodes with more connections to the seed list than expected by chance. For visualization, the resultant network obtained using CPDB was exported to yEd (https://www.yworks.com/products/yed; downloaded on August 1st, 2020).

### 2.5 Validation

For validation, we used the KM-plotter tool with two cohorts re-normalized with the Györffy dataset as part of a later compendium created by the same team. Specifically, we used the E-MBAT-365 (Guedj *et al*., 2012; Rème *et al*., 2013) and GSE20685 (Kao *et al*., 2011).

## 3 Results

### 3.1 Using Student’s t-test to rank genes by their association with 5-year RFS

Pre-processing and selection of the Györffy dataset resulted in a matrix with 13126 genes and 1519 individuals. Dividing this set by 5-year RFS resulted in two subsets: 689 patients who recurred within 5 years, and 830 patients who did not.

To look for genes that are associated with recurrence, we apply a slightly modified version of the common approach for microarray feature selection. According to the common approach, all the genes that show a statistically significant difference in their expression between these two patient groups, using the t-test (after multiple testing corrections) should be considered. However, this approach may not be suitable here since strikingly 67.8% of the genes (8901 of 13126 genes) are significantly differentially expressed between the patient groups after FDR correction. Because of the large number of differentially expressed genes, we repeated the analysis using the limma library to avoid potential p-value inflation due to multiple testing. Using limma we found 67.8% of the genes (9026 of 13126) to be significantly differentially expressed (see Supplementary Table S1 for the t-test and limma results for each gene). Moreover, a very high correlation was found between limma p-values and t-test p-values (r = 0.99, p-value < 2.2e-16). Thus, for the remaining analysis, we used the computationally simpler t-test.

To avoid working with such a large gene set, we slightly modified the regular approach by using the p-value of genes to rank them in decreasing order of significance. It is worth noting that since the degrees of freedom of all the genes are the same, the p-value of genes is proportional to their absolute t-statistic. After ranking genes by p-value, we chose the top 1% of all genes to define the “top 131 genes list”. The cutoff of 1% was chosen arbitrarily to ensure that all the chosen genes are highly significant (even after applying FDR cut-off <= 0.01) while the chosen set is sufficiently large for subsequent bioinformatics analyses. The distribution of absolute t-statistic values across all genes as well as the expression values distribution of the top-ranking 10 genes is summarized in Figure 1. From the analysis of the distribution of absolute t-test value, 3 gene groups emerge (Figure 1A). The first has a low absolute t-statistic (i.e. expected to have higher p-values), fitting a normal distribution with a mean of 0.9 and a standard deviation of 0.6. The second group of genes has a medium t-statistic and fits a normal distribution (3.4±1.4, mean and standard deviation). Finally, a group of genes is observed with a high t-statistic distribution (7.8±2.5). Tempting as it is to offer an explanation for this 3-modal distribution, it is risky to suggest a different role for genes from the 3 groups: the t-statistic considers not only the difference in means but also the variation. Since many biological and technical factors affect the variation in measured gene expression, it is hard to interpret the difference in t-statistic between genes.

**Figure 1.**
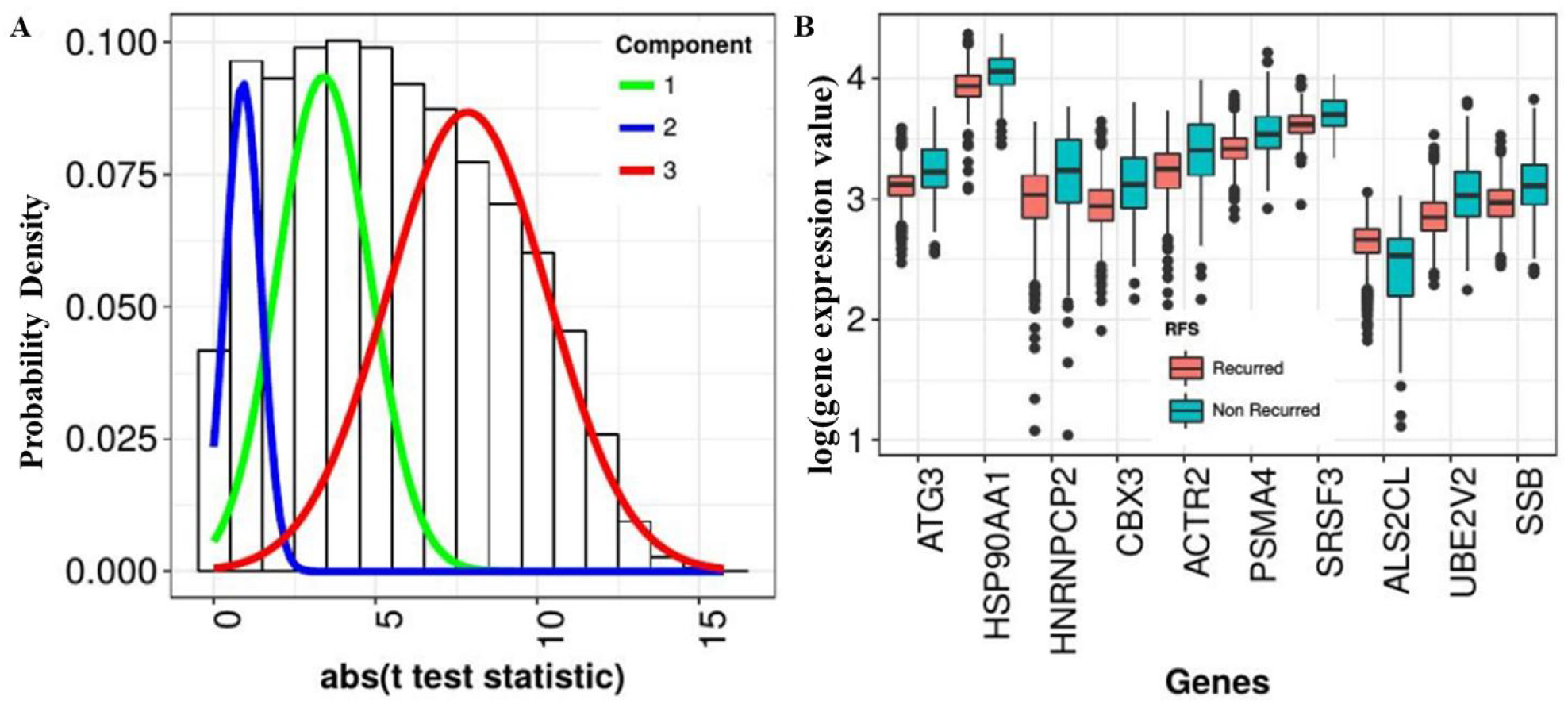
The t-statistic and gene expression differences associated with 5-year Recurrence Free Survival (RFS) in the Györffy breast cancer microarray dataset. The 1519 patients in the Györffy data set were grouped into those with a fast (<5year) vs slow (>5year) cancer recurrence. The difference in gene expression level between the two groups was computed using t-test for all the 13,126 genes and the adjusted p-value was used for ranking the genes. (A) A histogram (black-bordered bars) and probability density plot (tri-modal distribution in colors) of the t-test statistic values obtained using a normal best fit model. (B) A comparison of the top 10 ranked gene’s expression distribution for fast (in blue) and slow/no-recurrence cases (in red). The distribution of expression values (natural-log transformed) is summarized in a box-and-whiskers plot showing the minimum, median, maximum, and outliers (black dots).

To further validate the value of the t-test results for choosing informative genes in the context of this work, we summarize the expression of the 10 top-ranking genes in terms of p-values. For each gene, expression is compared between the fast (left) and slow/no recurrence (right) groups (Figure 1B). Clear differences in the expression values of each gene can be easily discerned, regardless of overall expression level: ALS2CL and HSP90AA1 differ 50-fold in their overall expression, yet both are in the top 10 genes in terms of the significance of the difference between groups. Interestingly, 9 out of the 10 genes most different between fast and slow recurrence cases have higher expression in the slow/no recurrence tumors: the only exception is ALS2CL, which is lower in slow/no recurring tumors.

### 3.2 HNF4A as a key gene in BC recurrence

We next tested if the top 131 genes list is enriched in specific pathways using EnrichR over-representation analysis. We identified several pathways as significantly enriched when considering REACTOME pathways (see Table 1 for the top 10, and Supplementary Table S2 for all pathways). Among the enriched pathways, multiple immune-related pathways emerge, such as “innate immunity” (with an adjusted p-value of 0.004); several cell-cycle related pathways are also enriched, such as “M phase” (adjusted p-value=0.004) or “role of GTSE1 in G2/M progression after G2 checkpoint” (adjusted p-value=0.004); “Cell Cycle Mitotic” (adjusted p-value=0.004) and several signal transduction pathways such as “VEGFA-VEGFR2 pathway” (adjusted p-value=0.004), “CLEC7A (Dectin-1) signaling” (adjusted p-value=0.004) and “Activation of NF-kappaB in B cells” (adjusted p-value=0.004).

**Table 1:** The over-represented pathways significantly enriched within the top-1% ranked genes in the Györffy breast cancer microarray dataset. The top 1% (n=131) of all the ranked genes in the Györffy data that differentiate between fast (<5year) vs slow (>5year) breast cancer recurrence was used to perform over-representation analysis using EnrichR online tool. The over-represented KEGG (top) and REACTOME (bottom) pathways are identified.

| Term/pathway name | Overlap | Adjusted p-value* |
| --- | --- | --- |
| **Source: KEGG** |  |  |
| Proteasome Homo sapiens | 4/44 | 0.033310 |
| **Source: REACTOME (top 10)** |  |  |
| Immune system homo sapiens | 26/1547 | 0.003637 |
| M-phase homo sapiens | 10/268 | 0.003637 |
| The role of GTSE1 in G2/M progression after G2 checkpoint homo sapiens | 5/59 | 0.004088 |
| CLEC7A (Dectin-1) signaling homo sapiens | 6/99 | 0.004088 |
| VEGFA-VEGFR2 pathway homo sapiens | 10/320 | 0.004088 |
| Cell Cycle, Mitotic homo sapiens | 12/462 | 0.004088 |
| Signaling by VEGF homo sapiens | 10/328 | 0.004088 |
| Activation of NF-kappaB in b cells homo sapiens | 5/66 | 0.004088 |
| Innate immune system homo sapiens | 16/807 | 0.004088 |
| Cell cycle homo sapiens | 13/566 | 0.004088 |

### 3.3 Induced network analysis suggests a role for HNF4A in fast BC recurrence

The overrepresentation analysis indicated that the top 131 genes list constitute a non-random group of genes. However, it is hard to understand, from this analysis, which genes are related to BC recurrence and how: genes may participate in more than one pathway, and pathway response may be secondary to the biological process underlying the propensity for rapid recurrence. We thus decided to augment the enrichment analysis by applying the induced network modules analysis algorithm from CPDB, using the top 131 genes list as input.

Induced network analysis with CPDB with the default cutoff for inferred nodes (i.e. 20 standard deviations above the average) identifies a tightly connected network (Figure 2a). In fact, 87 out of 131 genes list in the top 131 genes list are connected in this network, again suggesting that the top 131 gene list is non-random, and suggesting that these genes are involved in one or more biological functions that pertain to recurrence propensity.

**Figure 2.**
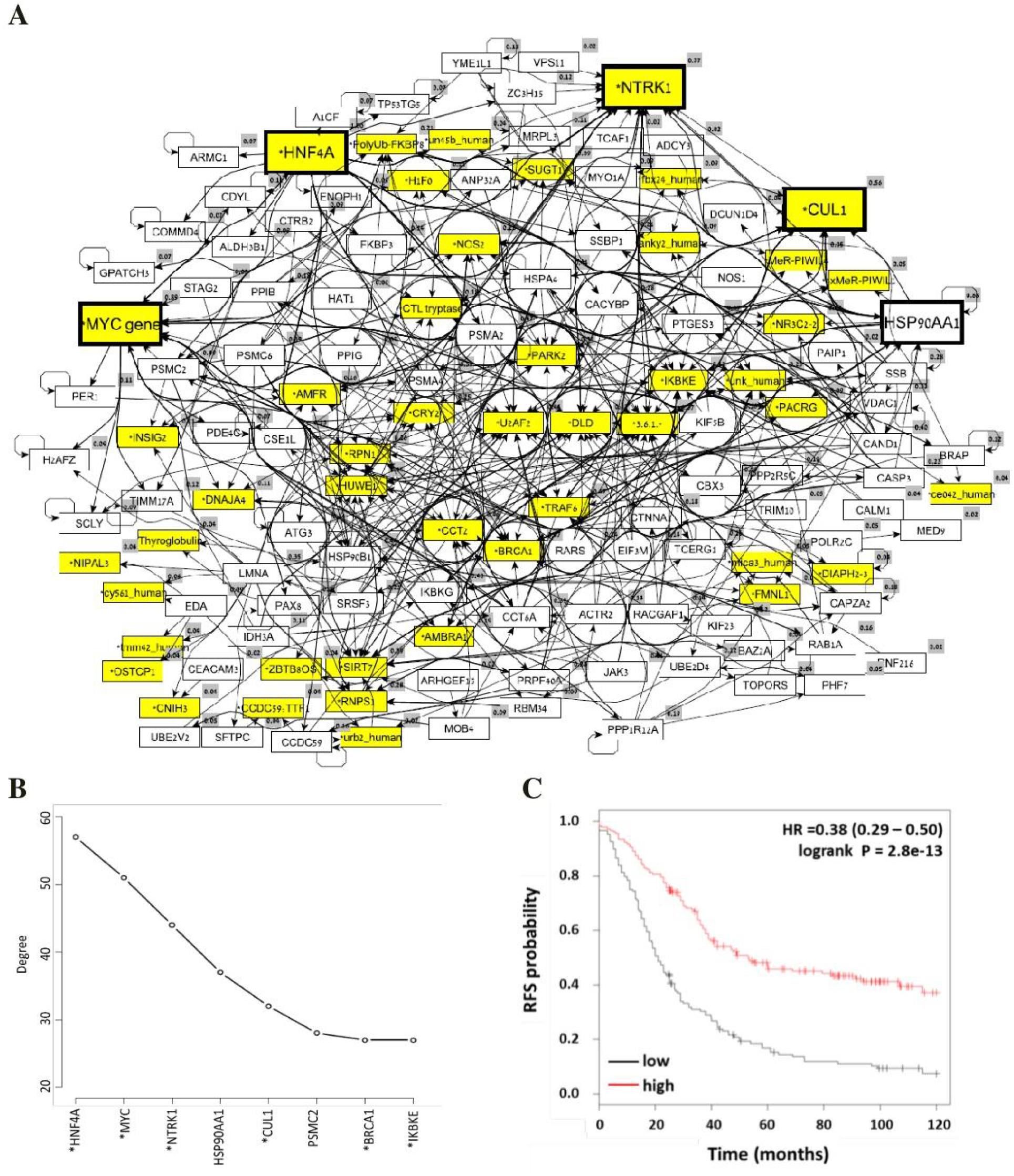
Induced network module analysis of top 1% t-test ranked genes from Györffy breast cancer microarray dataset. The top 1% (n=131) of all the ranked genes in the Györffy data with the highest p-value that differentiate between fast (<5year) vs slow (>5year) cancer recurrence was passed to the induced network module of the CPDB online tool. The resulting gene connection network (A) was further extended to inferred regulators or interactors (marked with ‘*’ preceding the name) not present in the input gene list using a binomial proportion test and a connectivity cutoff. Larger boxes indicate the 5 most connected genes. (B) The 8 genes with the most number of connections were identified. Induced genes (i.e. genes not in the top 1% list) are indicated with a star (*). (C) The significant association of HNF4A and recurrence free survival (RFS) is computed using KM-plotter tool and Györffy dataset. The recurrence free survival probability was plotted against time for patients with high expression(red) of the gene and for patients with low expression(black) of the gene.

In addition to revealing a network of connections between genes in the top 131 genes list, the induced network analysis infers highly connected genes, many of which are not actually in the top 131 genes list. To concentrate on the pivotal inferred genes, we first identified hubs: when the distribution of gene degrees (i.e. the number of connections per gene) is examined, a threshold of 30 connections emerges as a cutoff defining hubs (Figure 2b). Five genes are identified as hubs in the induced network with this cutoff: HNF4A, MYC, NTRK1, HSP90AA1, and CUL1. Two interesting observations are worth considering here. First, HSP90AA1 is the only hub that is in the top 131 gene list; the other 4 are inferred nodes, i.e., they have more connections to genes in the network than expected by chance; high expression of HSP90AA1 has been shown to be a marker of poor prognosis in HER2 negative BC (Cheng *et al*., 2012). The second observation is that 3 out of the 5 hub genes have been reported to be oncogenes (MYC and NTRK1) or tumor suppressors (HNF4A). The latter is the most connected gene in the induced network, with 57 connections. The remainder of this work thus concentrates on HNF4A.

HNF4A is a promiscuous transcription factor, with recorded interactions with 1878 genes in the Györffy dataset (according to InnateDB (Breuer *et al*., 2013)). If HNF4A activity is related to time to recurrence in BC, we expect HNF4A targets to occur more commonly in the top 131 genes list. Analysis of the induced network supports this prediction: of the 131 genes in the top 131 genes list, 87 genes are included in the connected network. Of these, 26 are connected to HNF4A, compared to the 12 expected by chance (χ2 = 17.2, p = 0.00003). In other words, HFN4A targets are 2-fold more likely to be in the top 131 genes list than expected by chance and statistical analysis suggests this enrichment is unlikely to be a coincidence.

HNF4A was identified not because it is in the top 131 gene list but because it was highly connected to genes in the RFS-related list. This raises the question of whether HNF4A expression is associated with RFS. To test this question, we used KM plotter to compare RFS in HNF4A high and HNF4A low patients (Figure 2c) using probe id 230772_at. This analysis indicates that HNF4A expression is strongly associated with RFS time, with an HR of 0.38 (90% CI: 0.29-0.5; p-value = 3·10-13, log-rank test).

Since HNF4A is a transcriptionally regulated transcription factor (TF) that differs in expression level between fast and slow/no recurring patients, it is reasonable to expect its targets to also be differentially expressed between these groups. However, since TFs may serve both as activators and repressors, some genes can be over-expressed while others may be suppressed by HNF4A activity. To overcome this complication, the correlation in expression between any two HNF4A targets was calculated. The resulting correlation matrix (Figure 3) reveals two groups of closely correlated genes (r>0.6, p<0.05). The first group includes GPATCH3, APOC4, COMMD4, CTNNA1, A1CF, MYO1A, ALDH3B1, PAX8 and positively correlates with HNF4A itself. These genes inversely correlate with the other group of HNF4A targets which are also all positively correlated to each other, including TCERG1, FKBP3, CCDC59, HSP90B1, BRAP, PTGES3, VDAC1, ARMC1, CCT6A, MRPL3, SSBP1, TIMM17A, PSMA2, CAPZA2, CBX3, ATG3, CACYBP and ZC3H15. This suggests that the first group includes genes activated by HNF4A, and the second contains genes that it represses.

**Figure 3.**
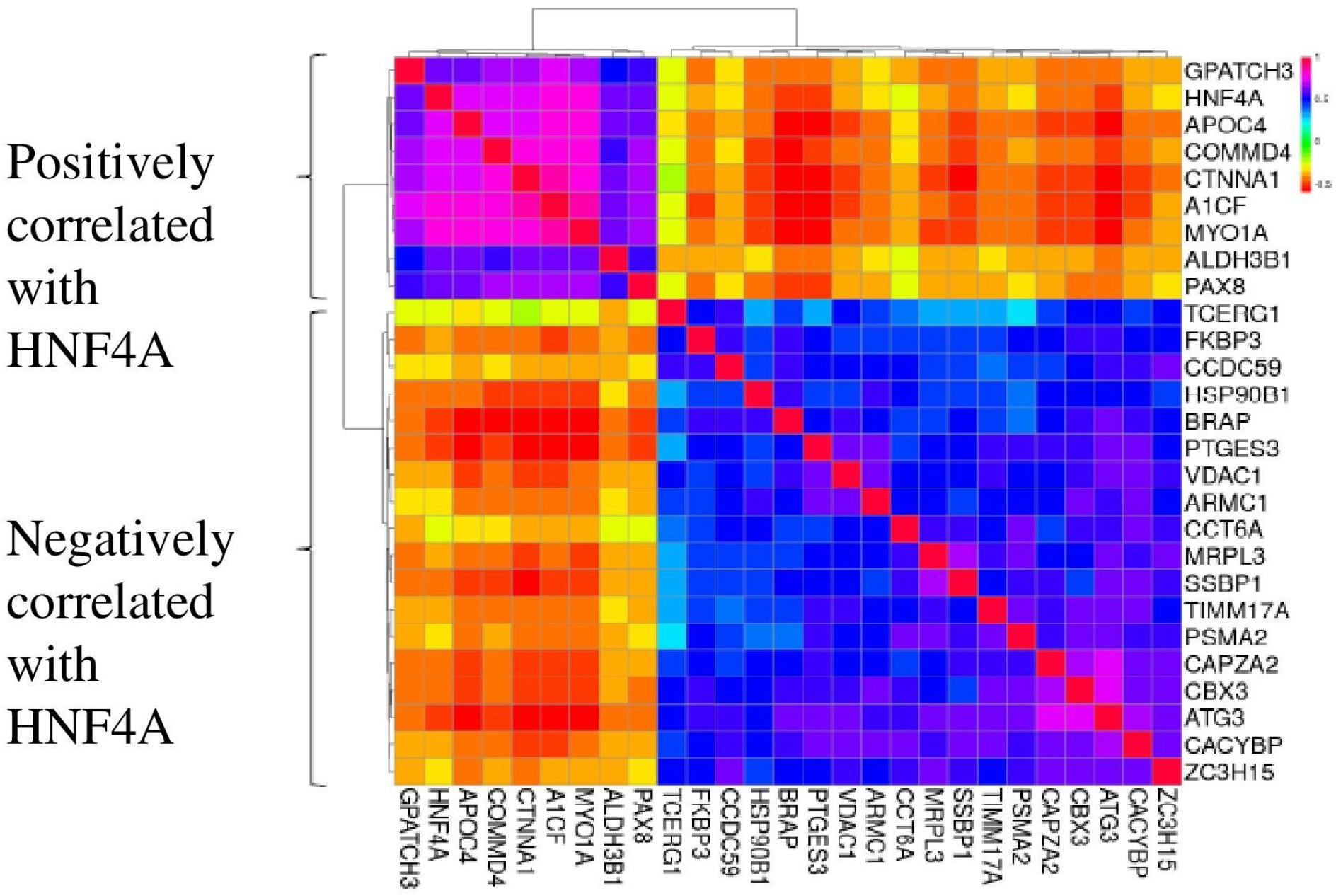
Gene expression correlation heatmap of HNF4A and HNF4A target genes from the Györffy breast cancer microarray dataset. The top 1% (n=131) of all the ranked genes in the Györffy data with the highest p-value was used to create an induced network module. The induced network module analysis identified 26 target genes of HNF4A. A gene expression correlation heatmap was constructed for HNF4A and its 26 target genes. The genes that are positively correlated & activated by HNF4A and negatively correlated and repressed by HNF4A in the Györffy dataset are shown.

Since these are all genes from the top 131 genes list, their association with RFS time (using log-rank test) cannot be considered evidence of their association with recurrence. However, the choice of the top 131 genes list was based on a mixed cohort (involving ER+ and ER-patients for which ER status is known, N=905 and 243 correspondingly). Since the recurrence of the ER+ subtype is slower than ER-, the predictive value of HNF4A targets in separated sub-cohorts, containing ER+ or ER-patients, is not trivial. Among the HNF4A-regulated genes in the top 131 genes list and HNF4A itself, 23/27 were found to be associated with RFS concordantly between ER+ and ER-patients, which is much more than expected by chance (Table 2, p=3·10-4, chi-square test). Strikingly, the Hazard Ratio (HR) was concordant between ER+ and ER-patients even when the log-rank test was not significant. In other words, even if the difference in RFS time was not statistically significant, the impact of high or low expression was almost always associated with a concordant impact on RFS time. Since the lack of significance can be easily explained by the relatively small size of the sub-cohorts, we further investigated HR concordance rather than significance in the log-rank test for the remainder of this work.

**Table 2.**
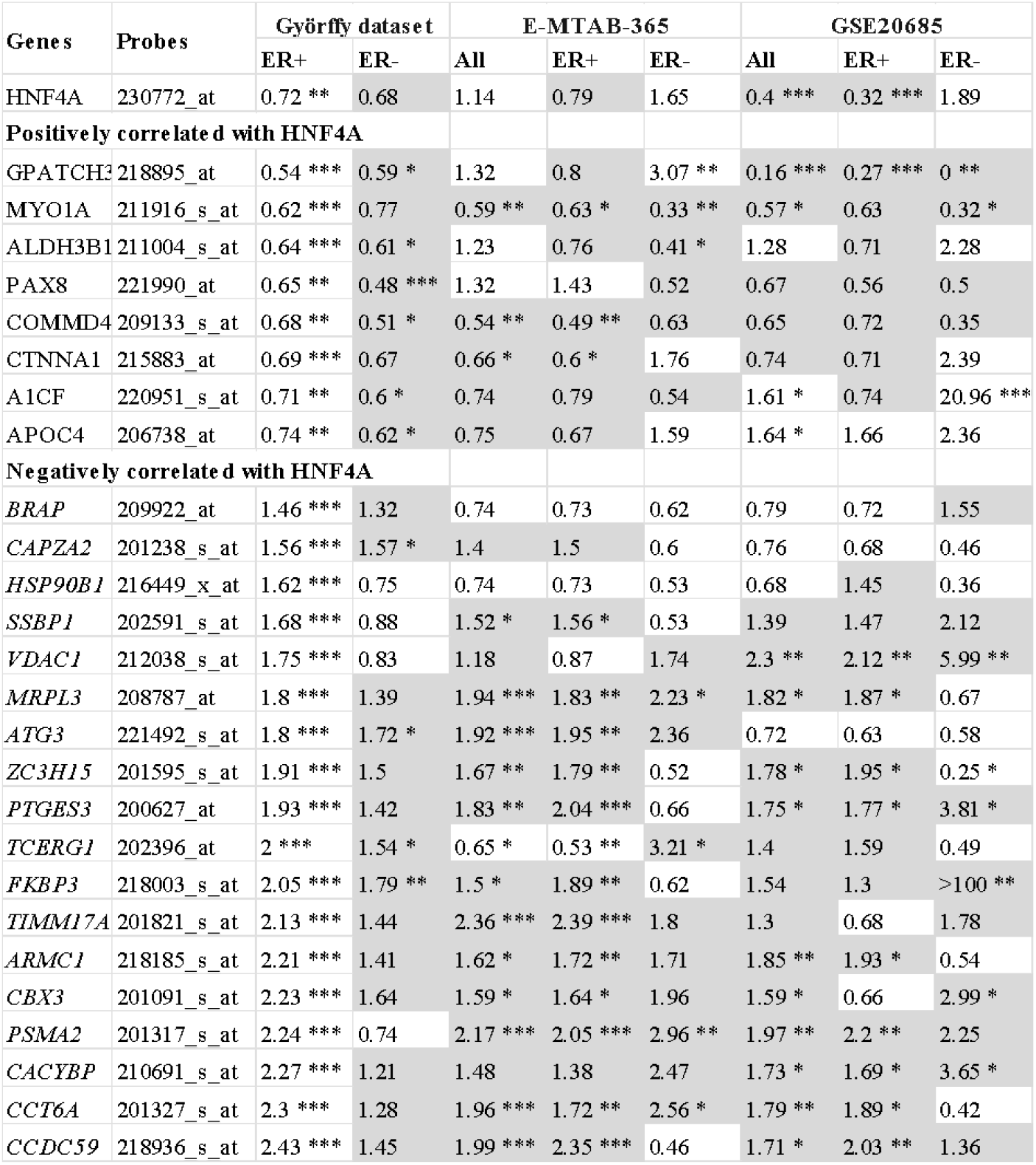
A strong concordance in the direction of HNF4A expression for its top 1% targets with recurrence free survival (RFS). The association of HNF4A and its target genes to recurrence free survival time is validated using KM plotter for Györffy dataset and two independent breast cancer microarray data cohorts (E-MTAB-365 and GSE20685). Patients were divided by estrogen receptor (ER) status (+/−). For the independent cohorts, only patients who received endocrine and chemotherapy treatment were considered. For each gene, the significance in association between gene expression status and recurrence free survival is computed using KM plotter tool. The value for Hazard Ratio (HR) is numerically indicated, and significance is indicated with stars (’***’ indicates p≤0.001; ‘**’ indicates p≤0.01 and ‘**’ indicates p≤0.05, using unadjusted log-rank test results from KM plotter). For each HNF4A target, the direction of its correlation with HNF4A is also indicated (positively or negatively correlated). Concordance with the ER+ subset of the Györffy dataset is indicated by dark shading.

### 3.4 Validation of HNF4A and the expression of its target in two independent breast cancer cohorts

To finally validate the predictive power of HNF4A expression and its targets in the top 131 genes list, their association with RFS was analyzed with KMplotter considering independent cohorts, not included in the Györffy dataset. In particular, we considered two datasets, E-MBAT-365 and GSE20685, which had the highest number of samples (N = 426 and N = 327 respectively). To further test the clinical utility and to ensure homogeneity, we chose only cases that received endocrine- and chemotherapy. High expression of HNF4A is associated with longer RFS in ER+ patients from both validation sets, although this association was significant only for GSE20685 (Table 2).

When the targets of HNF4A from the top 131 genes list are considered in the validation set, a mixed picture emerges with a tendency toward concordance with the findings of the Györffy dataset. In the E-MBAT-365 ER+ sub-cohort, 22/27 genes are concordant with the ER+ sub-cohort of the Györffy dataset. Moreover, significant association with RFS time is observed for 16 genes; all but one of these is concordant in the direction of association between HNF4A expression and RFS time. The ER+ sub-cohort of the GSE20685 validation set has high (21/27 genes) concordant with the Györffy dataset ER+ sub-cohort. Again, the significance of association increases the chances of concordance: 11 of the 11 significant associations found for this sub-cohort are concordant with the Györffy dataset ER+ sub-cohort.

It is worth noting that much weaker concordance is observed when comparing ER-cases from the validation sets and the Györffy dataset ER+ sub-cohort. Only 15 and 14 (of 27) genes are concordant for E-MBAT-365 and GSE20685, correspondingly. Significant associations are also less common (7 and 9 for E-MBAT-365 and GSE20685, correspondingly). Interestingly, a strong yet discordant association is observed for one HNF4A target in the ER-sub-cohorts of the validation sets: A1CF, for which high expression is associated with longer RFS (HR<1) for most cohorts, has a significant but inverse correlation with RFS in All or ER-patients from the GSE20685 cohort (p<0.001, HR>20). It is worth noting here that we only considered endocrine and chemotherapy-treated patients, and the RFS we observe reflects the response of ER-patients to such treatment.

HNF4A is a linoleic acid receptor. It is not unlikely that RFS improvement can be achieved by modulating linoleic acid levels. If this is indeed the case, we expect enzymes involved in linoleic acid metabolism to also be associated with RFS (Chiang, 2009). Testing CERS2 and 4 suggests this is indeed the case (Figure 4): while weak or no association is observed for most CERS, a significant association is found with CERS2 and CERS4, which best works with C18 carbon chains; Linoleic acids are a C18 chain (Sassa and Kihara, 2014).

**Figure 4.**
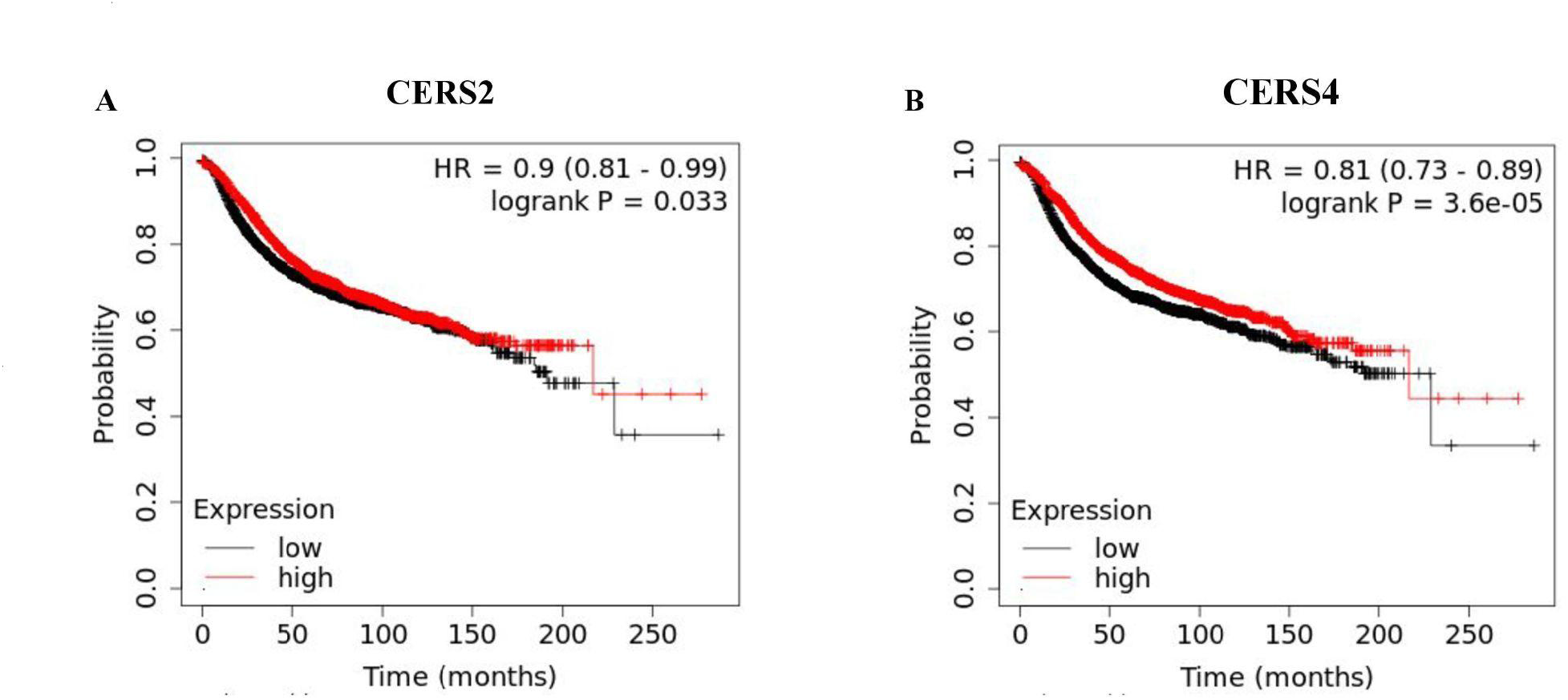
The association between CERS2 and CERS4 with recurrence free survival time. An induced network module analysis of the top 1% (n=131) of all the ranked genes in the Györffy data set identified HNF4A, a linoleic acid receptor, to be the most central gene associated with breast cancer recurrence free survival. The significance in association between linoleic acid metabolizing enzymes CERS2 & CERS4 to recurrence free survival (RFS) was further conducted using KMplotter tool. The recurrence free survival probability was plotted against time for patients with high expression(red) of the gene and for patients with low expression(black) of the gene. (A) The significant association of CERS and RFS (B) The significant association of CERS4 and RFS.

## 4 Discussion

In this work we apply a pipeline slightly modified from the standard expression analysis to gain new insights into the mechanism of BC recurrence. Beginning with a traditional method, we use a compendium of BC expression data to contrast genes expression between patients with fast (<5 years) and slow/no recurrence. We continue with the customary Student’s t-test to compare the significance of the difference in expression between the two groups, but instead of taking all the differentially expressed genes, we choose the top 1% of all the ranked genes which are significant, i.e. 131 genes, as the “top 131 genes list”. This is followed by a standard overrepresentation analysis of this list of genes. This analysis identifies immune-related pathways, including innate immunity pathways, cell-cycle related pathways, and signal transduction networks.

We started our analysis by mapping 22,215 probes to their corresponding genes using gprofiler, followed by choosing a probe with the lowest p-value for every gene resulting in 13126 genes expression value. We used probe choice rather than probe summation to extract a statistic per gene when genes had multiple probes. We chose an approach that maximizes the differential expression potential of each gene. This means that any other probe choice/summation method would have to achieve the same or less significant p-values for each gene compared to the method we used, which in turn could alter the way genes are ranked. However, even if probe summation/choice methods can introduce noise into gene ranking, they cannot specifically enrich top ranking genes with targets of HNF4A. We thus conclude that even if it is interesting to test other, more sophisticated methods to calculate differential expression, in the scope of this work our simplistic approach is sufficiently accurate to define a biologically meaningful group of genes.

In this work, we used the top 1% of genes because too many genes had a significant difference in expression between the groups (67.8% of the genes had a statistically significant p-value, an FDR threshold of 0.01). This can be explained in multiple ways. Residual uncorrected batch effects may inflate differences in expression between batches. However, we have used data that has been re-normalized to reduce batch effects (as explained in the methods sections). Moreover, since there is no overlap between batches and recurrence (i.e. All batches contain both slow/no fast recurrence patients) batch effects cannot fully explain the large proportion of differentially expressed genes. Lack or imperfect multiple testing correction could in principle also contribute to apparent differential expression. However, we used multiple testing corrections with a stringent cut-off (q-value <=0.01). Moreover, limma also found a large proportion of genes that are differentially expressed using the same stringent multiple testing corrected significance cut-off (Supplemental figure S2). Our probe selection method could also inflate the significance of difference since we chose for each gene the probe with the most significant p-value. However, this and all other sources of inflation are expected to randomly assign significance to genes. Yet, enrichment analysis reveals of the top 131 differentially expressed genes suggesting this is not a random list. Thus we cannot rule out the possibility that some of the differential genes that we report as expressed actually non-variable but seem so due to one or more artifacts. Yet non-variable genes are not likely to be dominant in the top 131 genes list. in the top 131 genes list.

To augment overrepresentation analysis, we use the Induced Network Module Analysis tool of CPDB to identify network complexes that biologically function together from our top 131 gene list. This tool takes a list of genes, extracts gene-gene connections between them (and to other genes/proteins outside the list) from several databases such as InnateDB, and builds a network of interconnected genes. The tool goes further to infer additional genes/proteins that are not in the original list but are more connected to proteins in the list than expected by chance. For this work, the induced network module analysis tool identifies proteins/genes that are more tightly connected to the top 131 genes list than expected by chance. This approach identifies 2 known cancer genes, MYC and NTRK1 as inferred hubs, confirming the validity of the analysis (Figure 2). It also infers HNF4A as the major hub connected to many genes in the top 131 genes list, as well as 2 more hub genes CUL1 and HSP90AA1.

Three lines of evidence in our results support the hypothesis that HNF4A expression level is associated with RFS time: (1) it has 2-fold more targets in the top 131 genes list than expected by chance; (2) its expression is significantly associated with RFS time, with good concordance in HR direction between ER+ and ER-sub cohorts, as well as in validation cohorts that were not used to choose HNF4A (Table 2); and (3) the expression of the 26 HNF4A targets that are in the top 131 are also associated with RFS, with better concordance in the direction of RFS differences than expected by chance.

The results we report with the Györffy dataset are largely reproducible in smaller yet independent cohorts. When E-MTAB-365 and GSE20685 are considered, even when strict patient selection criteria are applied to ensure homogeneity in treatment, the results are largely concordant with those found in the original discovery dataset (Table 2). The direction of the association between gene expression and RFS time was largely concordant with the ER+ sub-cohort of the Györffy dataset for both validation sets, and the tendency toward concordance was much stronger for genes for which the difference in RFS was statistically significant. One exception is worth noting here: A1CF is discordant and highly singingly associated with RFS in the GSE20685 ER-cohort, while it’s concordant in the ER+ sub-cohort. However, the size of this validation cohort, the complexity of the interactions between gene expression and response to therapy, and the possibility that A1CF expression could be just a marker for another transcription factor make it impossible to draw firm conclusions from the exceptional behavior of this gene.

HNF4A is a transcription factor and a nuclear hormone receptor with functions including bile acid synthesis, drug metabolism, differentiation, cell proliferation, immune response, and apoptosis (Bolotin *et al*., 2010; Walesky and Apte, 2015). In the context of cancer, its expression is shown to be associated with cancer risk in the kidney, liver, and colon cancers (Grigo *et al*., 2008; Babeu *et al*., 2009; Walesky and Apte, 2015). Low expression of HNF4A has been associated with metastasis, invasion, and epithelial-to-mesenchymal transition (EMT) in neuroblastoma and colon cancers (Alotaibi *et al*., 2015; Xiang *et al*., 2015; Jucá *et al*., 2017). A recent review on nuclear receptors as therapeutic targets in breast cancer postulates that HNF4A is a tumor-suppressor and that developing therapeutic strategies that activate it or increase its expression would be useful (Garattini *et al*., 2016). It has also been shown to be associated with RFS in ER+ and ER-patients and is proposed to be involved in tumor proliferation through the regulation of cellular metabolism in breast cancer (Aesoy, Clyne and Chand, 2015). However, the regulatory role of HNF4A in breast cancer recurrence is not understood (Huber-Keener *et al*., 2012; Aesoy, Clyne and Chand, 2015).

It is very tempting to propose that HNF4A expression plays a causative role in BC recurrence. It is not hard to find ways by which a promiscuous transcription factor such as HNF4A, directly or indirectly regulates proteins that affect BC recurrence risk. HNF4A controls, for example, the expression of AURKA (Cerami *et al*., 2011) one of the proliferation markers in the Oncotype Dx assay(Genomic Health) BC recurrence score (Paik *et al*., 2004; Cronin *et al*., 2007), suggesting a role for HNF4A in regulating cell proliferation. This is only one example, and further investigation is required to test if and via which pathways HNF4A expression impacts recurrence likelihood. This possibility is of great clinical interest: should it be shown that HNF4A expression is protective against recurrence; new avenues for intervention will be opened. HNF4A is a nuclear hormone receptor of linoleic acid. Could avoiding dietary linoleic acid, the ligand of HNF4A, increase RFS or even prevent recurrence? There is already some evidence that connects linoleic to BC metastases (Arab *et al*., 2016), raising the possibility that potent regulators of HNF4A could be effective measures in delaying or preventing BC recurrence.

As mentioned above, causation cannot be inferred from correlation alone. It is not unlikely that some other processes or pathways are active or inactive in fast-recurring patients, and that these processes/pathways also affect HNF4A expression levels. If this is the case, HNF4A may be a marker of recurrence risk rather than a driver. In this case, it would be interesting to check if it adds on top of the existing recurrence score(s). Since HNF4A is a nuclear hormone receptor, it is possible that it would help resolve the recurrence risk in some patient subpopulations, thus increasing the accuracy of recurrence risk assessment. It should be noted that although such an improvement is yet to be directly demonstrated, our finding that HNF4A expression is associated with RFS even in ER-negative BC patients (Table 2), for whom Oncotype Dx is not currently helpful, supports this possibility. Nevertheless, this hypothesis with a compendium of patients for whom (1) recurrence risk was estimated with the existing recurrence scores e.g. Oncotype Dx (Genomic Health); (2) HNF4A expression was measured; and (3) patients were followed sufficiently to record RFS. Such data are currently unavailable.

The HNF4A expression pattern could also be indicative of response treatment, rather than reflect the natural progression of the disease. HNF4A regulates potent drug-metabolizing enzymes, such as CYP2C9 (Chiang, 2009), and has been proposed to play a major role in tamoxifen metabolism (Jernström *et al*., 2009). Since almost all the patients in the compendium used for this work were treated, the RFS used in our analysis reflects the response to the standard of care. It is not impossible that the difference between fast and slow/no recurrence is due to drug metabolism or other drug-response related differences between the patients. This model is supported, at least in part, by the association of HNF4A expression levels with tamoxifen resistance in a cell-line based analysis (Huber-Keener *et al*., 2012).

We are well aware that it is customary to divide patients by their clinical subtypes. However, we did not divide the patients in the hope to find genes that are associated with recurrence across subtypes. Our results suggest that this hope has been realistic: the association of HNF4A, which we found without dividing the patients by subtypes, was found to be associated with recurrence in ER+ only and ER-only patients. We did not further divide ER-negative patients into triple negative and HER2+ since the latter groups were too small for statistical power. Nevertheless, our approach and the very large dataset we have been using may have resulted in a generic and robust recurrence-associated gene.

To conclude, in this work, we bring evidence that HNF4A expression/activity is associated with time to recurrence in BC. Further experiments are required to test if HNF4A is causatively linked to recurrence, directly or indirectly, via treatment response, or if it is a marker of some recurrence-related process. If it is causative, our results open new therapeutic avenues; if it is a marker; it could add information on top of existing markers/scores for estimating recurrence risk. In either case, because HNF4A is such a major transcription factor, our work suggests that further investigation of its association with BC recurrence is required.

## 6 Author contributions

RVV and ER conceived the study. RVV, NEA, RH, and ER performed the data analysis. ER supervised the study. RVV and ER drafted the manuscript. All authors read and approved the final manuscript.

## 9 Data Availability Statement

The code and datasets analyzed for this study can be found in the Github repository [https://github.com/vvrahul11/BRRP-GED].

## 13 Supplementary Figures

**Supplementary figure 1:**
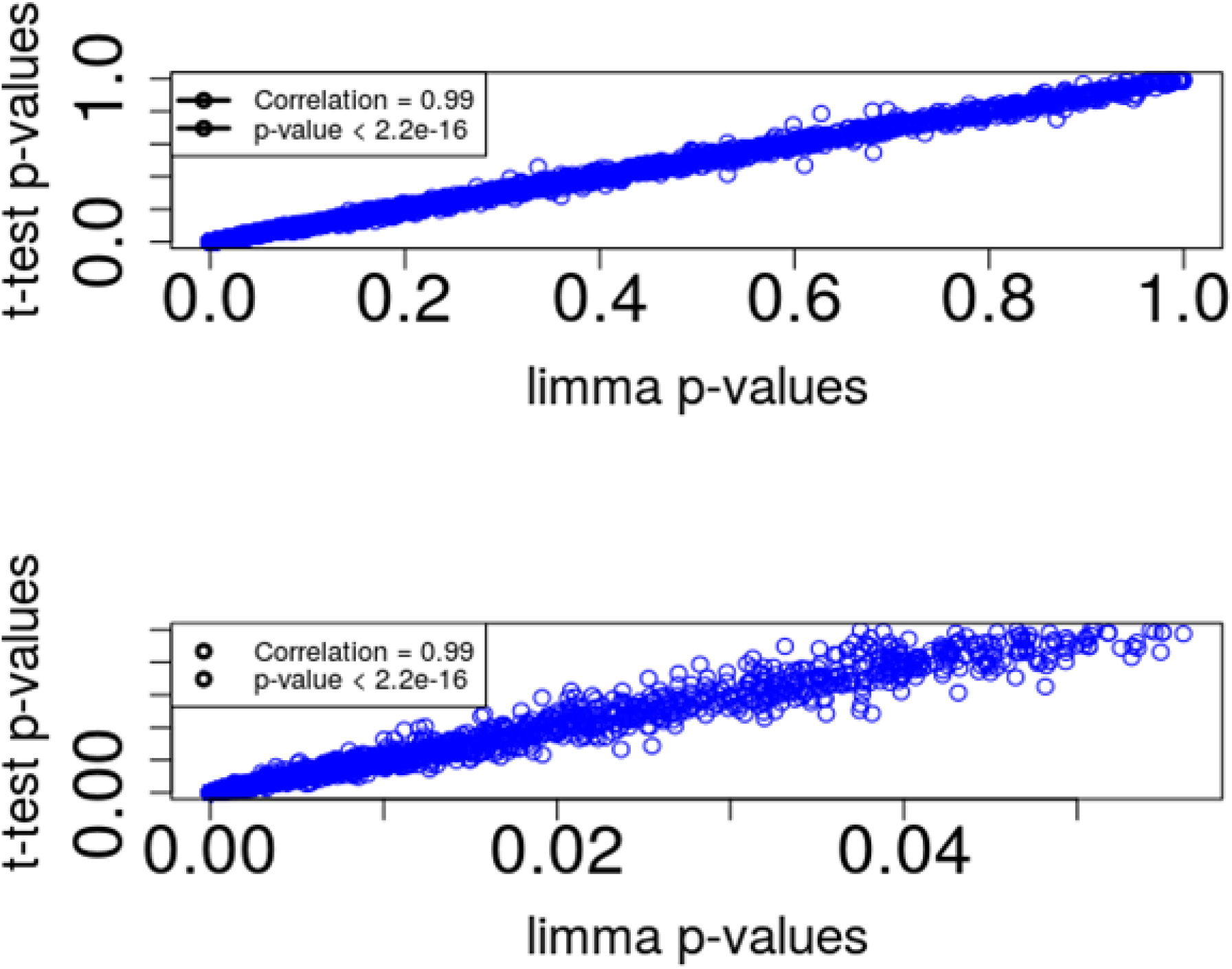
The correlation between t-test and limma for selected probes. Representative probes were selected for each gene as described in materials and methods (briefly: the probe with the most significant t-test statistic was chosen for each gene). The p-value for every probe was compared between limma (horizontal axis) and t-test (vertical axis). (A) The entire p-value range. (B) only probes with t-test derived p-value of 0.05 or lower are shown.

## References

Aesoy, R., Clyne, C.D. and Chand, A.L. (2015) ‘Insights into Orphan Nuclear Receptors as Prognostic Markers and Novel Therapeutic Targets for Breast Cancer’, Frontiers in Endocrinology, 6. Available at: https://www.frontiersin.org/articles/10.3389/fendo.2015.00115 (Accessed: 19 September 2022).

Alotaibi, H. et al. (2015) ‘Enhancer cooperativity as a novel mechanism underlying the transcriptional regulation of E-cadherin during mesenchymal to epithelial transition’, Biochimica Et Biophysica Acta, 1849(6), pp. 731–742. Available at: 10.1016/j.bbagrm.2015.01.005.

Arab, A. et al. (2016) ‘The effects of conjugated linoleic acids on breast cancer: A systematic review’, Advanced Biomedical Research, 5, p. 115. Available at: 10.4103/2277-9175.185573.

Babeu, J.-P. et al. (2009) ‘Hepatocyte nuclear factor 4alpha contributes to an intestinal epithelial phenotype in vitro and plays a partial role in mouse intestinal epithelium differentiation’, American Journal of Physiology. Gastrointestinal and Liver Physiology, 297(1), pp. G124–134. Available at: 10.1152/ajpgi.90690.2008.

Benaglia, T. et al. (2010) ‘mixtools: An R Package for Analyzing Mixture Models’, Journal of Statistical Software, 32, pp. 1–29. Available at: 10.18637/jss.v032.i06.

Bolotin, E. et al. (2010) ‘Integrated approach for the identification of human hepatocyte nuclear factor 4alpha target genes using protein binding microarrays’, Hepatology (Baltimore, Md.), 51(2), pp. 642–653. Available at: 10.1002/hep.23357.

Braun, S. et al. (2003) ‘Evaluation of bone marrow in breast cancer patients: prediction of clinical outcome and response to therapy’, Breast (Edinburgh, Scotland), 12(6), pp. 397–404. Available at: 10.1016/s0960-9776(03)00143-7.

Breuer, K. et al. (2013) ‘InnateDB: systems biology of innate immunity and beyond--recent updates and continuing curation’, Nucleic Acids Research, 41(Database issue), pp. D1228–1233. Available at: 10.1093/nar/gks1147.

Carter, C.L., Allen, C. and Henson, D.E. (1989) ‘Relation of tumor size, lymph node status, and survival in 24,740 breast cancer cases’, Cancer, 63(1), pp. 181–187. Available at: 10.1002/1097-0142(19890101)63:1<181::aid-cncr2820630129>3.0.co;2-h.

Cerami, E.G. et al. (2011) ‘Pathway Commons, a web resource for biological pathway data’, Nucleic Acids Research, 39(Database issue), pp. D685–690. Available at: 10.1093/nar/gkq1039.

Chen, E.Y. et al. (2013) ‘Enrichr: interactive and collaborative HTML5 gene list enrichment analysis tool’, BMC Bioinformatics, 14(1), p. 128. Available at: 10.1186/1471-2105-14-128.

Cheng, Q. et al. (2012) ‘Amplification and high-level expression of heat shock protein 90 marks aggressive phenotypes of human epidermal growth factor receptor 2 negative breast cancer’, Breast cancer research: BCR, 14(2), p. R62. Available at: 10.1186/bcr3168.

Chiang, J.Y.L. (2009) ‘Hepatocyte nuclear factor 4alpha regulation of bile acid and drug metabolism’, Expert Opinion on Drug Metabolism & Toxicology, 5(2), pp. 137–147. Available at: 10.1517/17425250802707342.

Coley, H.M. (2008) ‘Mechanisms and strategies to overcome chemotherapy resistance in metastatic breast cancer’, Cancer Treatment Reviews, 34(4), pp. 378–390. Available at: 10.1016/j.ctrv.2008.01.007.

Cronin, M. et al. (2007) ‘Analytical validation of the Oncotype DX genomic diagnostic test for recurrence prognosis and therapeutic response prediction in node-negative, estrogen receptor-positive breast cancer’, Clinical Chemistry, 53(6), pp. 1084–1091. Available at: 10.1373/clinchem.2006.076497.

Demicheli, R. et al. (2004) ‘Menopausal status dependence of the timing of breast cancer recurrence after surgical removal of the primary tumour’, Breast cancer research: BCR, 6(6), pp. R689–696. Available at: 10.1186/bcr937.

Dessau, R.B. and Pipper, C.B. (2008) ‘[“R”--project for statistical computing]’, Ugeskrift for Laeger, 170(5), pp. 328–330.

Fisher, B. et al. (2002) ‘Twenty-year follow-up of a randomized trial comparing total mastectomy, lumpectomy, and lumpectomy plus irradiation for the treatment of invasive breast cancer’, The New England Journal of Medicine, 347(16), pp. 1233–1241. Available at: 10.1056/NEJMoa022152.

Garattini, E. et al. (2016) ‘Lipid-sensors, enigmatic-orphan and orphan nuclear receptors as therapeutic targets in breast-cancer’, Oncotarget, 7(27), pp. 42661–42682. Available at: 10.18632/oncotarget.7410.

Grigo, K. et al. (2008) ‘HNF4 alpha orchestrates a set of 14 genes to down-regulate cell proliferation in kidney cells’, Biological Chemistry, 389(2), pp. 179–187. Available at: 10.1515/BC.2008.011.

Guedj, M. et al. (2012) ‘A refined molecular taxonomy of breast cancer’, Oncogene, 31(9), pp. 1196–1206. Available at: 10.1038/onc.2011.301.

Györffy, B. et al. (2010) ‘An online survival analysis tool to rapidly assess the effect of 22,277 genes on breast cancer prognosis using microarray data of 1,809 patients’, Breast Cancer Research and Treatment, 123(3), pp. 725–731. Available at: 10.1007/s10549-009-0674-9.

H, W. (2016) ‘ggplot2: Elegant Graphics for Data Analysis. Springer-Verlag New York’. Available at: http://ggplot2.org.

Huber-Keener, K.J. et al. (2012) ‘Differential gene expression in tamoxifen-resistant breast cancer cells revealed by a new analytical model of RNA-Seq data’, PloS One, 7(7), p. e41333. Available at: 10.1371/journal.pone.0041333.

Irizarry, R.A., Gautier, L. and Cope, L.M. (2003) ‘An R Package for Analyses of Affymetrix Oligonucleotide Arrays’, in G. Parmigiani et al. (eds) The Analysis of Gene Expression Data: Methods and Software. New York, NY: Springer (Statistics for Biology and Health), pp. 102–119. Available at: 10.1007/0-387-21679-0_4.

Jafari, P. and Azuaje, F. (2006) ‘An assessment of recently published gene expression data analyses: reporting experimental design and statistical factors’, BMC Medical Informatics and Decision Making, 6, p. 27. Available at: 10.1186/1472-6947-6-27.

Jernström, H. et al. (2009) ‘CYP2C8 and CYP2C9 polymorphisms in relation to tumour characteristics and early breast cancer related events among 652 breast cancer patients’, British Journal of Cancer, 101(11), pp. 1817–1823. Available at: 10.1038/sj.bjc.6605428.

Jucá, P.C. de F.C., et al. (2017) ‘HNF4A expression as a potential diagnostic tool to discriminate primary gastric cancer from breast cancer metastasis in a Brazilian cohort’, Diagnostic Pathology, 12(1), p. 43. Available at: 10.1186/s13000-017-0635-2.

Kamburov, A. et al. (2009) ‘ConsensusPathDB—a database for integrating human functional interaction networks’, Nucleic Acids Research, 37(Database issue), pp. D623–D628. Available at: 10.1093/nar/gkn698.

Kao, K.-J. et al. (2011) ‘Correlation of microarray-based breast cancer molecular subtypes and clinical outcomes: implications for treatment optimization’, BMC Cancer, 11(1), p. 143. Available at: 10.1186/1471-2407-11-143.

Kolde, R. (2019) ‘pheatmap: Pretty Heatmaps’. Available at: https://CRAN.R-project.org/package=pheatmap (Accessed: 30 November 2022).

Kuleshov, M.V. et al. (2016) ‘Enrichr: a comprehensive gene set enrichment analysis web server 2016 update’, Nucleic Acids Research, 44(W1), pp. W90–97. Available at: 10.1093/nar/gkw377.

Lánczky, A. et al. (2016) ‘miRpower: a web-tool to validate survival-associated miRNAs utilizing expression data from 2178 breast cancer patients’, Breast Cancer Research and Treatment, 160(3), pp. 439–446. Available at: 10.1007/s10549-016-4013-7.

MP, C., et al. (2007) Cancer Incidence in Five Continents Volume IX. Available at: https://publications.iarc.fr/Book-And-Report-Series/Iarc-Scientific-Publications/Cancer-Incidence-In-Five-Continents-Volume-IX-2007 (Accessed: 18 September 2022).

Paik, S. et al. (2004) ‘A multigene assay to predict recurrence of tamoxifen-treated, node-negative breast cancer’, The New England Journal of Medicine, 351(27), pp. 2817–2826. Available at: 10.1056/NEJMoa041588.

Reimand, J. et al. (2016) ‘g:Profiler-a web server for functional interpretation of gene lists (2016 update)’, Nucleic Acids Research, 44(W1), pp. W83–89. Available at: 10.1093/nar/gkw199.

Rème, T. et al. (2013) ‘Modeling risk stratification in human cancer’, Bioinformatics (Oxford, England), 29(9), pp. 1149–1157. Available at: 10.1093/bioinformatics/btt124.

Ritchie, M.E. et al. (2015) ‘limma powers differential expression analyses for RNA-sequencing and microarray studies’, Nucleic Acids Research, 43(7), p. e47. Available at: 10.1093/nar/gkv007.

Saphner, T., Tormey, D.C. and Gray, R. (1996) ‘Annual hazard rates of recurrence for breast cancer after primary therapy’, Journal of Clinical Oncology: Official Journal of the American Society of Clinical Oncology, 14(10), pp. 2738–2746. Available at: 10.1200/JCO.1996.14.10.2738.

Sassa, T. and Kihara, A. (2014) ‘Metabolism of Very Long-Chain Fatty Acids: Genes and Pathophysiology’, Biomolecules & Therapeutics, 22(2), pp. 83–92. Available at: 10.4062/biomolther.2014.017.

Therneau, T.M. (2020) ‘Survival: A Package for Survival Analysis in R.’ Available at: https://CRAN.R-project.org/package=survival.

Valagussa, P., Tancini, G. and Bonadonna, G. (1987) ‘Second malignancies after CMF for resectable breast cancer’, Journal of Clinical Oncology: Official Journal of the American Society of Clinical Oncology, 5(8), pp. 1138–1142. Available at: 10.1200/JCO.1987.5.8.1138.

van’t Veer, L.J. and Bernards, R. (2008) ‘Enabling personalized cancer medicine through analysis of gene-expression patterns’, Nature, 452(7187), pp. 564–570. Available at: 10.1038/nature06915.

Walesky, C. and Apte, U. (2015) ‘Role of Hepatocyte Nuclear Factor 4α (HNF4α) in Cell Proliferation and Cancer’, Gene Expression, 16(3), pp. 101–108. Available at: 10.3727/105221615X14181438356292.

Wilke, C.O. (2020) ‘cowplot: Streamlined Plot Theme and Plot Annotations for “ggplot2”’. Available at: https://CRAN.R-project.org/package=cowplot (Accessed: 30 November 2022).

Xiang, X. et al. (2015) ‘Hepatocyte nuclear factor 4 alpha promotes the invasion, metastasis and angiogenesis of neuroblastoma cells via targeting matrix metalloproteinase 14’, Cancer Letters, 359(2), pp. 187–197. Available at: 10.1016/j.canlet.2015.01.008.

Yuan, X. et al. (2009) ‘Identification of an Endogenous Ligand Bound to a Native Orphan Nuclear Receptor’, PLOS ONE, 4(5), p. e5609. Available at: 10.1371/journal.pone.0005609.

